# Regional and age-dependent Effects of Cortical Magnetic Stimulation on Unconstrained Reaching Behavior

**DOI:** 10.1101/2020.12.14.422725

**Authors:** M.A. Urbin, Jing Tian, Gina P. McKernan, Nick Kortzorg, Lore Wyers, Florian van Halewijck, Matthieu P. Boisgontier, Oron Levin, Stephan P. Swinnen, Ilse Jonkers, George F. Wittenberg

**Affiliations:** VA Pittsburgh Healthcare System, Pittsburgh, PA, USA; Department of Physical Medicine & Rehabilitation, University of Pittsburgh, Pittsburgh, PA, USA; Department of Neurology, University of Maryland, Baltimore, MD, USA, Maryland Exercise and Robotics Center of Excellence, Geriatrics Research Educational and Clinical Center, Department of Veterans Affairs, Baltimore, MD, USA Laboratory for Research on Arm Function and Therapy, Older Americans Independence Center, Departments of Neurology, Physical Therapy and Rehabilitation Science, and Medicine/Division of Gerontology and Geriatric Medicine, University of Maryland, Baltimore, MD, USA; Department of Kinesiology, Biomedical Sciences Group, KU Leuven, Leuven, Belgium; School of Rehabilitation Sciences, Faculty of Health Sciences, University of Ottawa, Ottawa, ON, Canada; Bruyère Research Institute, Ottawa, ON, Canada

**Keywords:** motor control, aging, goal-directed movement, transcranial magnetic stimulation, dorsal premotor cortex

## Abstract

**Background:** The specific and dynamic contributions of premotor and supplementary motor areas to reaching movements in aging humans are not well understood.

**Objective:** To better understand the role of cortical motor regions and age on the control of unconstrained reaches against gravity by neurologically intact, younger and older adults.

**Methods:** Double pulse transcranial magnetic stimulation (TMS) was applied at locations targeting primary motor cortex (M1), dorsal premotor area (PMA), supplementary motor area (SMA), or dorsolateral prefrontal cortex (DLPFC). Paired stimuli were delivered before or after a visual cue was presented to initiate self-paced right-handed reaches to one of three, vertically oriented target locations.

**Results:** Regional stimulation effects on movement amplitude were observed both early and late in the reach. PMA stimulation increased reach distance to a greater extent than M1, SMA, and DLPFC stimulation. M1 and PMA stimulation increased deviation from the straight-line path around the time of peak velocity to an extent that was greater than SMA and DLPFC stimulation. Cortical stimulation increased the time that elapsed after, but not before, peak velocity. Despite stronger effects of stimulation on reaches in the younger group, this group had shorter times to reach the target after reaching peak velocity.

**Conclusion:** These results provide support for a role of PMA in visually guided movement *after* movement initiation. For older subjects, the increased time to arrive at the target after peak velocity despite weaker stimulation effects suggests an age-related reduction in sensorimotor processing flexibility for online control of unconstrained reaching.

**Highlights:**

- Dorsal premotor area stimulation at any time during the reaction-time period and early reaching affected early reach kinematics at least as much as stimulation of primary motor cortex.
- Older individuals had more stimulation-related interference in the late components of reaching despite having less early effect of stimulation, suggesting a reduction in flexibility of dynamic motor control due to aging.
- The antigravity component of unconstrained reaching did not have special aspects for regional cortical effects of stimulation.

## Introduction

Reaching for an object in space forms the basis of many activities of daily living. Contemporary views of goal-directed reaching emphasize the involvement of online processes that make maximal use of somatosensory and visual feedback to control limb trajectory.^1^ It is thought that two types of online processes regulate early- and late-phase control of reaching, respectively. Impulse control occurs early in the movement and involves a comparison of actual limb kinematics to an internal representation of expectations about limb trajectory. Limb-target control occurs late in the movement and involves real-time error reduction based on the relative position of the limb and target. Despite traditional views that the initial, ballistic phase of the movement is entirely preprogrammed given temporal limitations on sensory processing,^2^ it has been shown that unremitting availability of visual and somatosensory feedback enables online control, even for rapid movements performed under severe time constraints (i.e. movement times <150 ms). For example, limb trajectories adapt to changes in target size and location that occur after movement onset.^3^ Such online adjustments occur late in the movement,^4,5^ without conscious awareness.^6,7^

Over a half century of animal and human work has attempted to dissociate the role of various cortical motor areas (among all brain areas) in the planning and execution of goal-directed limb movement. Different features of activation in primary motor cortex (M1) are linked to kinematic^8-11^ and kinetic^12,13^ movement properties. However, the unique contribution of supplementary motor area (SMA) and premotor area (PMA), which both interconnect with M1,^14–16^ remains unclear. SMA, in the medial portion of Brodmann area 6, is thought to be involved in movement preparation^17^ and control of internally generated movement.^18,19^ Dorsal and ventral PMA, located lateral to SMA but also within Brodmann area 6, are thought to be involved in segregated fronto-parietal circuits subserving different aspects of reach-and-grasp movements. Dorsal PMA is thought to be more involved in the reach and ventral PMA in the grasp. Dorsal PMA receives dense projections from the dorsal visual stream, which provides a basis for visually guided movement.^20^ Although a considerable body of animal^21–23^ and human^24–26^ evidence supports the hypothesis that dorsal PMA is critically involved in sensory guidance and planning of goal-directed movement, recent work in non-human primates shows that inactivation of dorsal PMA impairs internally-generated but not visually-guided movements.^27^ So while the dorsal and ventral PMA are interesting in the context of visually-guided reaches, we studied only dorsal PMA here and use “PMA” to mean “dorsal premotor area.” (Localization of ventral PMA in human is also controversial^28,29^.) Also, while the targets were visual, there was reduced visibility of their location during the reach itself, so participants had to rely on an internal memory of target location.

Whether the cortical circuitry mediating online control is impacted by age during unconstrained reaching behavior is also not known. Prior work has shown increased end-point error and variability of reaches in older subjects^30^ and age-related differences in the relative contribution of ballistic and corrective movements.^31^ In these and most other studies, the reach task was simplified by restricting the movement to a two-dimensional plane with antigravity support of the arm.^30,32–34^ Such a reductionist approach has been useful but leaves open the question of whether there is any special consideration to the problem of countering the effects of gravity when a multijoint limb is lifted and extended away from the trunk.^35^ Age-related declines are observed in muscle mass,^36^ neuromuscular transmission,^37^ and cortical function,^38^ that support the ability to reach against gravity and harness passive inertial torques to achieve the goal of the movement.

The purpose of this exploratory study was to better understand the unique contributions of cortical motor areas to the control of unconstrained visually guided reaching behavior in neurologically intact humans. An ancillary purpose was to gain insight into the influence of age on control processes mediated by these brain regions. To accomplish both objectives, double pulse transcranial magnetic stimulation (TMS) was directed to M1, PMA, SMA, and dorsolateral prefrontal cortex (DLPFC) while younger and older subjects performed self-paced reaches to one of three targets. Targets were arranged vertically to capture varying gravitational demands on reaching behavior. A 100-ms interstimulus interval for double pulse TMS was used in these experiments because it has been shown to mainly disrupt cortical processing.^39^ Therefore, kinematic differences among stimulation conditions were used to infer a role of the brain region with activity disrupted by stimulation on controlling unconstrained reaching behavior.

## Methods

### Participants

18 individuals participated in the study and were recruited into young (n=10, 6 males, 26.4 ± 6.8 years) and old (n=8, 6 males, 67.4 ± 3.1 years) groups. All subjects reported good health, with no history of neurological diseases and normal or corrected-to-normal vision. Subjects were right-hand dominant, as established by the Edinburgh inventory.^40^ Signed informed consent was obtained from each subject prior to testing in accordance with policies of the local ethics committee and Declaration of Helsinki.^41^

### Experimental setup and task

Subjects performed all reaches while seated in a chair with backrest support and a restraint system situated around the trunk to restrict its movement. An adjustable table with a visual stimulus presentation system was placed in front of the subject (Fig 1A). Three pairs of light emitting diodes (LEDs) represented locations of an upper target (UT), middle target (MT) and lower target (LT). Both the LT and UT were vertically separated 15 cm from the MT. The height and horizontal positions of the MT were aligned to the right shoulder. The distance from the target to the subject was set to 5 cm less than a fully extended reach to the MT. The starting position was marked on the table and in line with the targets, 3 cm lower than the LT and 15 cm from the target board.

**Figure 1.**
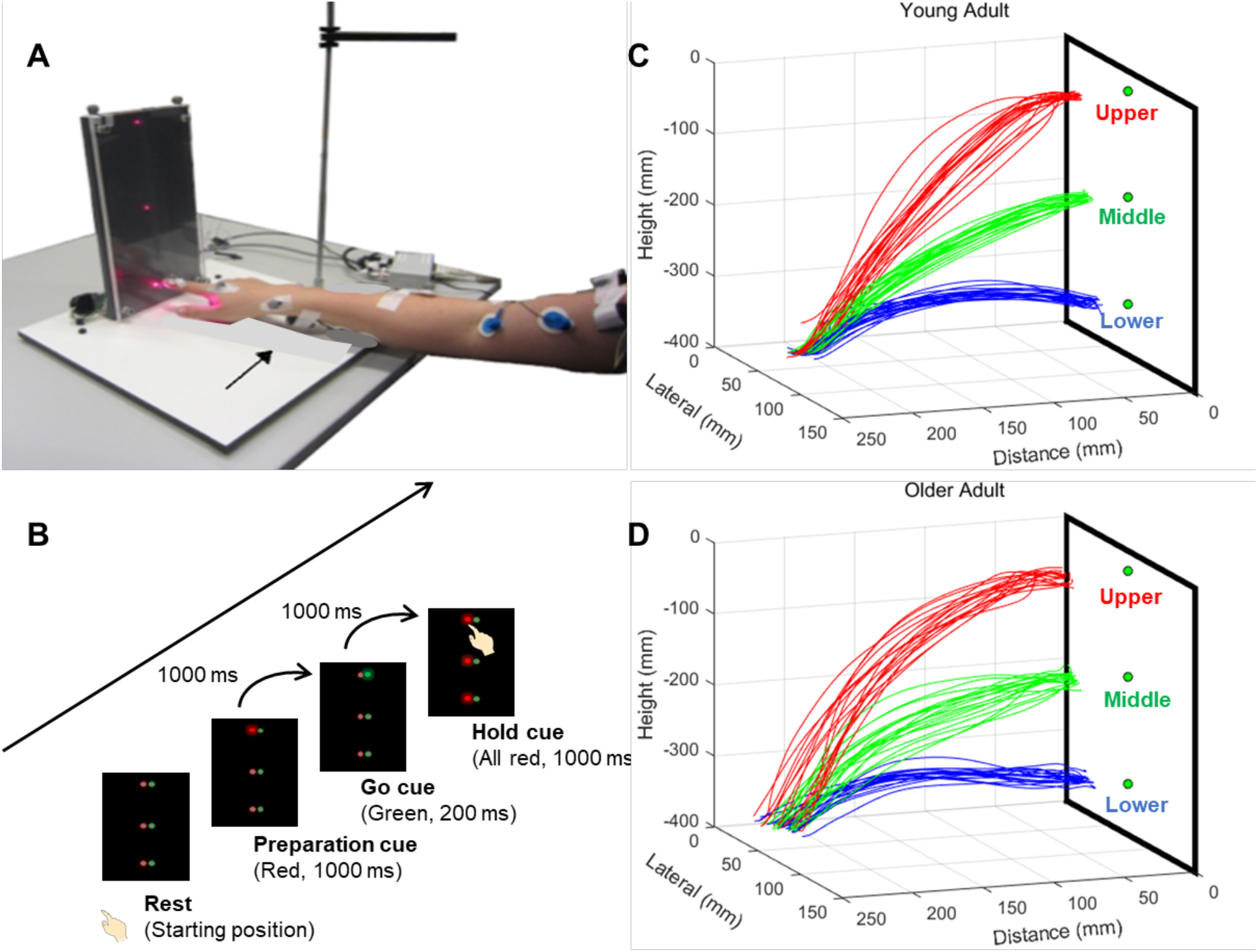
**A)** Visual stimulus presentation system and **B)** sequence of events in the unconstrained reaching task. Representative reach paths to the three target location from subjects in the **C)** younger and **D)** older groups.

Subjects were asked to find a comfortable sitting position with the right upper arm in a vertical, adducted position and elbow flexed approximately 90 degrees. The forearm was pronated with the hand resting on the table and the tip of index finger on the starting position. Each target location contained one red (left) and one green (right) LED separated by 1 cm (Fig 1B). Subjects were instructed to start at rest with the index finger on a starting position marker. Relaxation prior to movement initiation was stressed explicitly. First, one red LED illuminated for 1 s as a preparation cue, indicating the target for the forthcoming reach. A green LED illuminated for 200 ms after an additional 800-1200 ms random delay as a cue to initiate movement. All red LEDs illuminated 1 s after the movement initiation cue and remained illuminated for 1 s. Subjects were instructed to arrive at the target approximately 1 s after the initiation cue and maintain contact with the target until all red LEDs extinguished (i.e., for 1 s). The room light was dimmed, so visual feedback of the LED target and hand position was reduced but not eliminated.

### Experimental Procedure

A fixed pseudorandom sequence of 24 trial types was programmed in Turbo Pascal for DOS (Borland, Austin, TX) with 18 trials in which double pulse TMS was delivered at one of three timings and six trials with no stimulation. Reach target location varied across all 24 trials within a sequence, covering all combinations of target sequence and stimulation timing twice. Subjects performed a sequence of 24 practice reaches before two to three blocks using the programmed sequence were recorded with the coil at each of the four stimulation locations. This was done partly to help participants learn to approximate a 1-s movement duration. Paired stimuli separated by 100 ms were initiated at three different timings relative to the visual cue to initiate movement: 50 ms before cue (−50 ms), 100 ms after cue (+100 ms), and 250 ms after cue (+250 ms). Visual cues and double pulse TMS were controlled by the output of a parallel port of a DOS computer, connected to a custom-made trigger box.

### Transcranial Magnetic Stimulation

Double pulse TMS was delivered to left M1, left PMA, left SMA, and right DLPFC using a MagStim Rapid stimulator and standard 70 mm figure-eight coil. Inclusion of contralateral M1 was intended to serve as a positive control to dissociate the unique contributions of PMA and SMA (further explanation below). Although recent work has shown a role of DLPFC in disinhibiting contralateral M1 during *bilateral* movement preparation,^42,43^ ipsilateral DLPFC was intended to serve as a negative control with an expected negligible effect on unilateral reaching. DLPFC is also involved in working memory,^44^ but there was minimal demand to recall target location in the task paradigm described here. (For brevity, the “right” and “left” are omitted in most subsequent labels.)

A mapping procedure was administered in 10-20 space to localize the four brain regions stimulated during experimental procedures. Electromyography (EMG) was recorded (1 kHz) from the biceps brachii, long head of triceps brachii, anterior and posterior deltoids using bipolar surface electrodes (2-cm inter-electrode distance) affixed over each muscle belly. The stimulating coil was held tangentially to the scalp overlying M1 and angled 45 degrees laterally and posteriorly (MagStim 200^2^,70 mm figure-eight coil). The location that elicited the largest, peak-to-peak motor-evoked potential (MEP) in the right biceps brachii muscle from the mean of seven pulses was set as the optimal site. The same procedure was repeated for the tibialis anterior (TA) muscle with the stimulating coil at the midline of the scalp. During experimental procedures, M1 stimulation was applied at the biceps brachii optimal site, PMA stimulation was applied 2 cm anterior to this location, SMA stimulation was applied 3 cm anterior of the TA optimal site or 4 cm anterior to the vertex if a MEP could not be elicited, and DLPFC stimulation was applied 8 cm anterior to M1, mirrored to the hemisphere *ipsilateral to the reaching arm*.

Resting motor threshold (rMT) was determined as the minimum stimulator output needed to elicit a MEP with a peak-to-peak amplitude of > 50 μV in the biceps brachii muscle from 5 of 10 consecutive stimuli. Stimulator strength was set to 1.0x rMT for M1, 1.2x rMT for PMA, 1.5x rMT for SMA, and 0.8x rMT for DLPFC. Although the threshold of spinal motor neurons cannot be precisely controlled during a dynamic task, this stimulator output was intended to drive some degree of brief activation and more prolonged inhibition, with M1 stimulation as a positive control. Progressively higher stimulator outputs were used for PMA and SMA stimulation to increase the likelihood of measurable effects in absence of an efficient method to determine the intensity needed for such effects. Stimulator outputs were set at the lowest level possible to perturb the brain region targeted by stimulation, while limiting spread of activation to other brain regions. Suprathreshold stimulator outputs (i.e., 1.1x rMT) have also been used in previous work that aimed to perturb cortical areas during a motor task.^39^ This prior work established threshold based on visible twitch criteria. It is therefore likely that stimulator outputs used in the current study, which based rMT on the presence of 50-μV MEP amplitudes, were comparable to or below those used previously.

### Motion Capture

A ten-camera system (Vicon, Oxford Metrics, UK) in a fixed array along the ceiling edges of a motion capture laboratory, acquired the 3-dimensional location of reflective markers (100 Hz) affixed to anatomical landmarks on the arm (acromioclavicular joint, radial and ulnar styloid processes, humeral medial and lateral epicondyles, metacarpophalangeal joint and distal phalanx of the first finger) and trunk (C7 and T8 spinous processes, sternoclavicular joint, and xiphoid process). Three markers were placed on the stimulating coil, and four markers were placed on the head to track their positions. Figure 1 contains sample reach paths from a young (Fig. 1C) and old (Fig. 1D) subject that were reconstructed from motion capture data.

### Data Analysis

Kinematic data were processed offline using custom software developed in MATLAB™ (The Mathworks, Natick, MA). Data were passed through a low-pass (10 Hz), fourth-order, zero-lag Butterworth filter before further processing. Movement initiation and termination were defined as the instant when the velocity of the marker affixed to the metacarpal crossed above and below 5% of peak velocity in the vertical dimension, respectively. All trials were visually inspected to verify the validity of these criteria. Trials were removed from the analysis if movement was evident before the visual cue to initiate the reach, indicating a false start. Remaining trials were removed if reaction time or movement duration was >2 standard deviations from the mean, indicating missed cues or otherwise unusual kinematics.

Three positional parameters were calculated to characterize the amplitude of the reach path both early and late in the overall movement trajectory. The 3-dimensional reach vector was defined as the position of the index finger at each instant during the reach. Reach path magnitude was calculated by taking the square root of the sum of squares along each axis to characterize the extent of the reach path to the target. Reach path deviation was calculated by taking the difference between the reach path and a straight-line path to the target to characterize divergence of the reach path. Reach path curvature was calculated as the ratio between the overall length of the reach path and the distance between the position of the 3-dimensional reach vector at movement initiation and termination. Reach path magnitude and deviation were calculated early (i.e., 100 ms after movement initiation) and late (i.e., at the time of peak velocity and 50 ms thereafter) in the movement trajectory. The temporal structure of the reach was characterized by calculating the rise time and time after peak velocity.

EMG recordings from trials in the −50 ms condition were inspected to verify the incidence of MEPs in axial muscles integral to the reach (i.e., anterior deltoid, posterior deltoid, triceps longus, biceps brachii). The purpose was to determine whether and how frequently stimulation recruited spinal motor neurons. Recordings from the −50 ms condition were inspected, specifically, because both pre- and post-stimulation epochs were mostly free of background activation. All trials retained in kinematic analyses were used for this purpose. EMG recordings for these trials were rectified before waveform averaging trials corresponding to each target location. Mean EMG was calculated in the immediate 100 ms preceding stimulation onset. MEPs were detected if EMG increased 2 standard deviations above the mean of the prestimulus epoch and remained above this threshold for 15 ms consecutively, 10-30 ms after stimulation onset. The incidence of MEPs was expressed as a fraction of the total possible occurrences (3 targets x 18 subjects = 51 possible occurrences) for each muscle. MEP incidence was also calculated by target location.

### Statistical Analyses

Independent samples t-tests were used to test for differences between younger and older groups in age and rMT. The GLM procedure in SAS (Version 9.4) was used to perform overall F tests and the independent effects of brain region, target location, stimulation timing, and age group on kinematic parameters. Least square means and least square mean differences were computed for classification effects with multiple comparison adjustments for the p-values and confidence limits for the least square mean differences for individual parameters. We chose to model at the trial level to capture systematic trends in within-subject variability. The variability of human movement is well established and is likely exacerbated by the lack of constraint in the reaching task described here. Moreover, we anticipate that self-paced reaches would result in subjects more readily engaging online control processes to facilitate reaches to the target.

## Results

Subjects completed all research procedures within a single testing session. During that session an average of 65±9 trials (mean ± SD) were performed in the no-stimulation condition and at each stimulation timing for each subject. A subset of trials was removed for each condition based on the rejection criteria described above. The fraction removed in each condition were: no stimulation (14.9±7.5%), 50 ms before cue (8±8.5%), 100 ms after cue (7±9.1%), 250 ms after cue (6.5±9.2%). The younger averaged 26±7 years old years and older 67±3 years. There was no significant difference in rMT between younger (52.7±10.3%) and older (65.9±17.3%) groups (t_16_ = −1.9, *p*=0.084), although the older group had a trend towards a higher threshold, which is consistent with previous work^45^.

Prior to testing for kinematic differences, reaction times were examined in stimulation and no stimulation conditions to determine if there was an effect of stimulation. Such a *cueing* effect might bias movement kinematics under stimulation conditions due to earlier movement initiation. A mixed model on reaction time (F_95,4252_=10.39, *p*<0.001) revealed an effect of stimulation timing with −50 and +100 ms conditions producing significantly (all *p*<0.001) reduced reaction times relative to no-stimulation and 250-ms conditions (Fig. 2).

**Figure 2.**
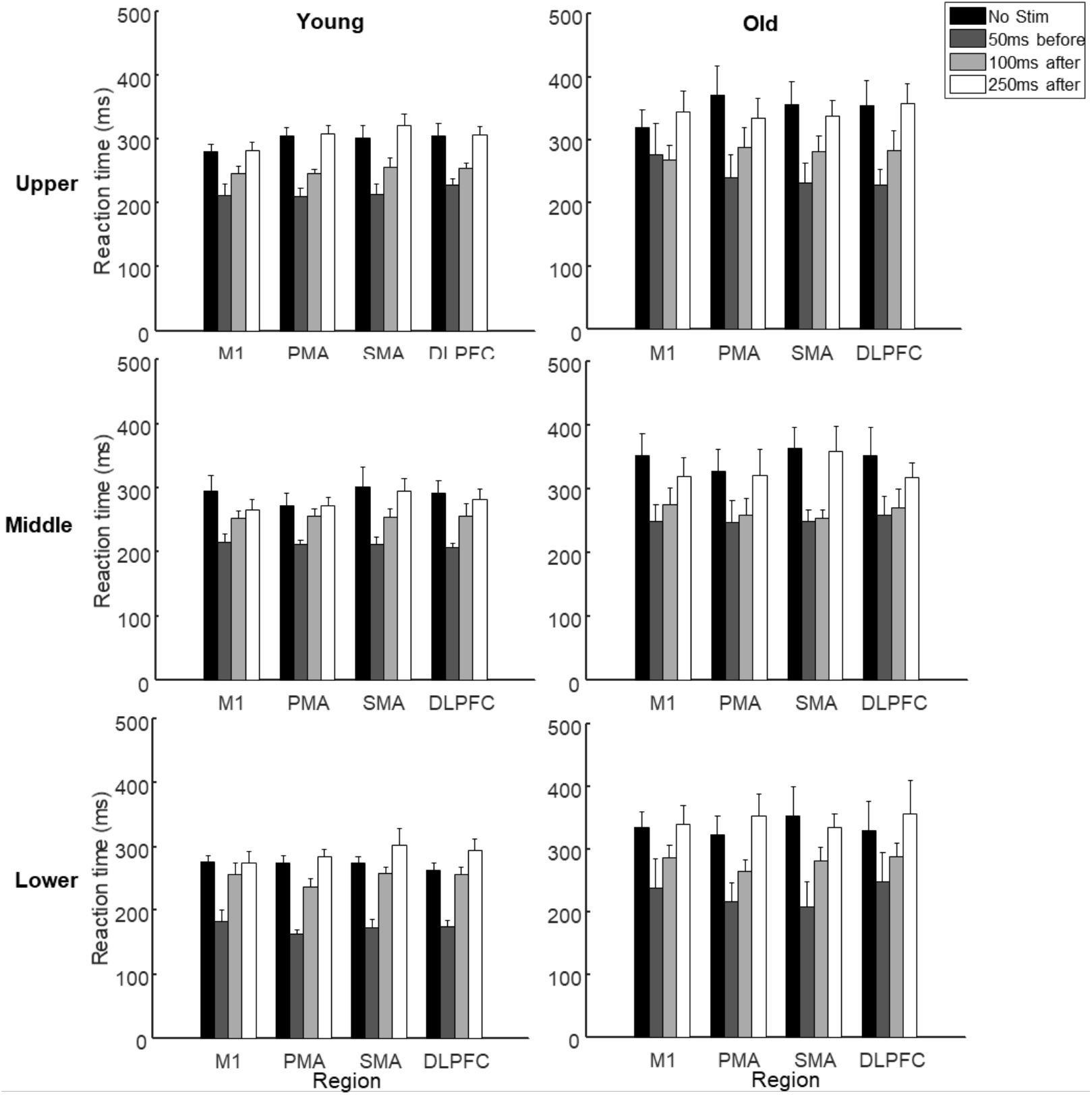
Group means of reaction time in younger (left column) and older (right column) subjects at each target location (rows) under no-stimulation and stimulation conditions. Note the reduced reaction time for both groups at each target location under each 50-ms before cue and 100-ms after cue stimulation conditions. (Error bars reflect the standard deviation).

Given evidence of a cueing effect, preliminary models excluded the no stimulation condition and focused on the regional effect of stimulation at all three timings. Models revealed a distinct pattern of regional, but not temporal, effects. PMA stimulation increased the magnitude of the reach path at 100 ms after movement onset to a greater extent than all other regions (M1, t= 4.452, *p*<.0001; SMA, t= 3.694, *p*=.0002; DLPFC, t= 8.603, *p*<.0001). Reach path magnitude at 100 ms after movement onset was also greater for M1 and SMA relative to DLPFC (t= 4.182, *p*<.0001 and t= 4.947, *p*<.0001, respectively). Mixed models for reach path deviation revealed that M1 and PMA stimulation increased deviation to an extent that was similar and greater than SMA and DLPFC stimulation throughout the reach, including 100 ms after movement onset (M1 > SMA, t= 3.46, *p*=.0005 and DLPFC, t= 2.992, *p*=.0028; PMA > SMA, t= 2.992, *p*=.0028 and DLPFC, t= 2.992, *p*=.0028), at the time of peak velocity (M1 > DLPFC, t= 2.992, *p*=.0028; PMA > SMA, t= 5.104, *p*<.0001 and DLPFC, t= 4.624, *p*<.0001), and 50 ms after the time of peak velocity (M1 > SMA, t= 2.32, *p*=.0272 and DLPFC, t= 4.07, *p*<.0001; PMA > SMA, t= 2.21, *p*=.0204 and DLPFC, t= 1.763, *p*=.078). The same pattern was observed for reach path curvature (M1 > SMA, t= 2.792, *p*=.0053 and DLPFC, t= 3.927, *p*<.001; PMA > SMA, t= 3.792, *p*=.0002 and DLPFC, t= 4.919, *p*<.0001).

In summary, preliminary statistical models revealed distinct patterns in regional effects of cortical stimulation. Since reaction time did not differ between no stimulation and 250-ms stimulation conditions, a final set of statistical models included only these conditions to avoid the confound of reaction time effects while also preserving the ability to observe pure regional effects of stimulation. Thus, all levels of brain region (4), target location (3), and age group (2) were retained, but only 2 levels of stimulation (no stimulation and 250-ms stimulation) were included in these models.

For reach path magnitude, an effect for brain region was only evident *early* in the movement trajectory (100 ms after movement onset (F_2,89_) =8.1, *p*<0.001, Fig 3). Exemplar reach path magnitudes with stimulation to each cortical region are shown in Figure 4. Similar to preliminary statistical models, reach path magnitude at 100 ms after movement onset was greater with PMA stimulation relative to all other regions (M1, t= 2.899, *p*=.0038, SMA, t= 3.6, *p*=.0003, and DLPFC, t= 6.162, *p*<.0001). M1 and SMA stimulation also produced a greater reach path magnitude than DLPFC (t= 3.175, *p*=.0002 and t= 2.997, *p*=.0031, respectively). Effects of stimulation on reach path magnitude were greater in the younger relative to the older group (*p*<0.001 and *p*<0.001, respectively), and a region by group interaction (*p*=0.04) was detected. There was a significant effect of target (*p*<0.001) and a target by group interaction (*p*<0.02), such that increased demands to counter the effects of gravity at successively higher target locations *decreased* reach path magnitude (UT<MT<LT) with the younger group exhibiting greater magnitudes.

**Figure 3.**
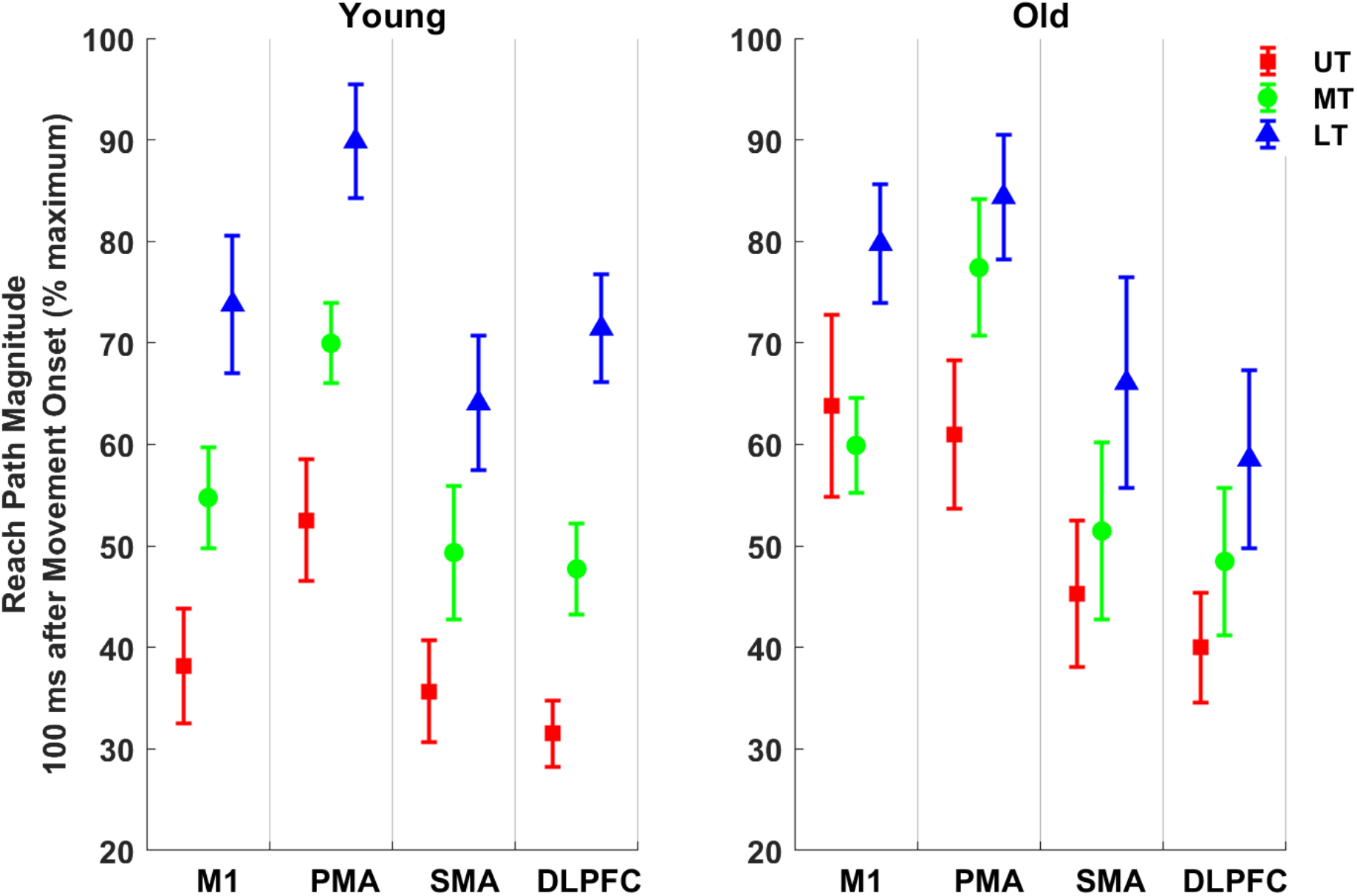
Trends in the regional effects of cortical stimulation on relative reach path magnitude 100 ms after movement onset in younger (left column) and older (right column) subjects at each target location under the +250-ms stimulation condition. Error bars represent standard error. Reach magnitudes across trials for all regions were normalized to the maximum magnitude from a given subject.

**Figure 4.**
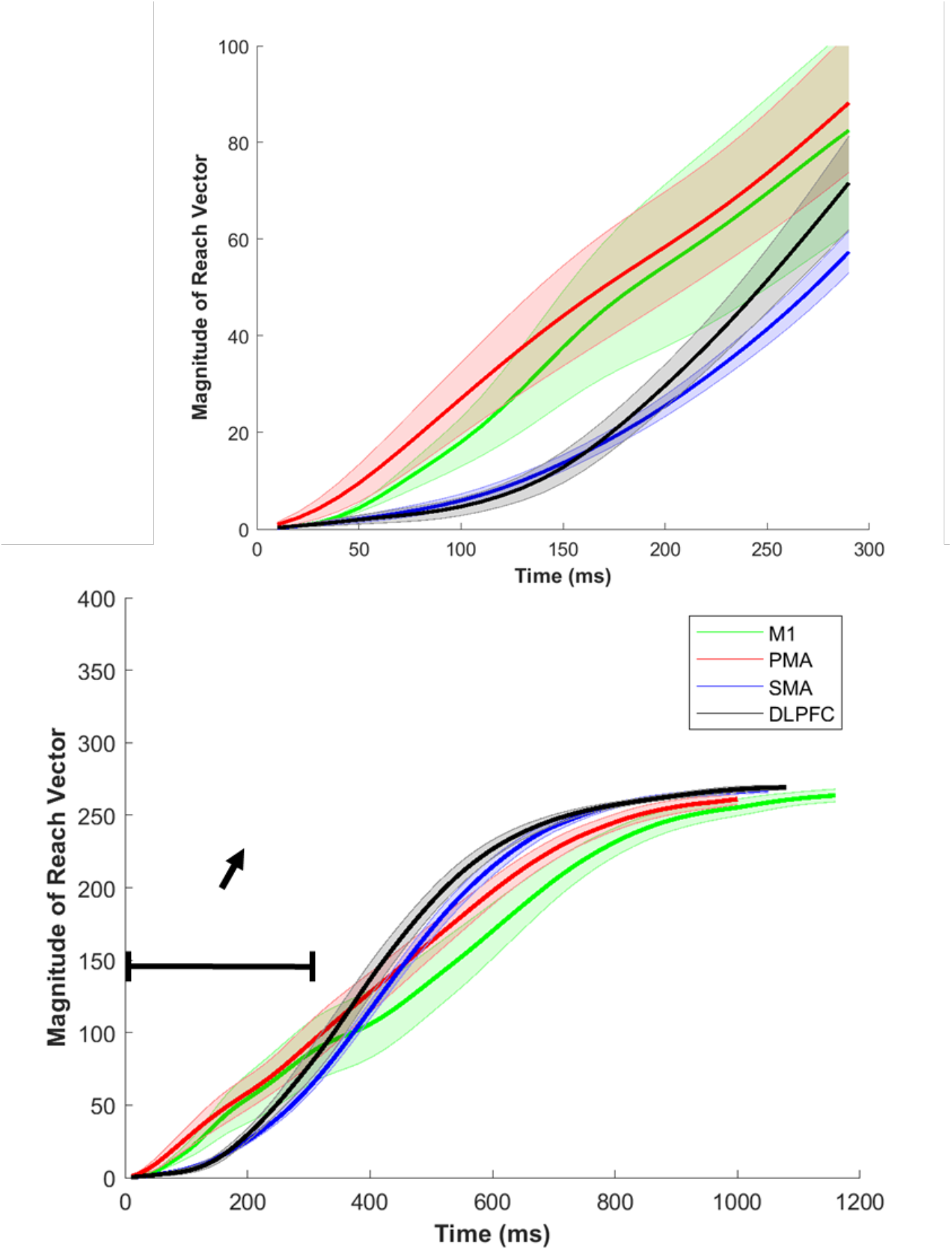
Exemplar reach path magnitude from a single subject under the 250-ms stimulation condition. The bottom plot illustrates magnitude of the reach vector over the entire reach, whereas, the top plot depicts only the first 300 ms of the reach to more clearly show regional differences detected at 100 ms after movement onset. Traces represent the mean of all trials when stimulation was delivered to a particular region, and shaded areas correspond to standard error.

For reach path deviation, there was reversal in trends such that regional effects were only evident *late* in the movement trajectory (at peak velocity: (F_2,89_) =15.11, *p*<0.001, Fig 5; 50 ms after peak velocity: (F_2,89_) =8.98, *p*<0.001, Fig 6). Exemplar reach path deviations with stimulation to each cortical region are shown in Figure 7. Just as preliminary models showed, reach path deviation was similar for M1 and PMA, with both regions increasing deviation beyond DLPFC and SMA stimulation at both the time of peak velocity (M1 > SMA, t= 2.694, *p*=.0071 and DLPFC, t= 2.423, *p*=.016; PMA > SMA, t= 3.428, *p*=.0006 and DLPFC, t= 3.161, *p*=.002) and 50 ms thereafter (M1 > SMA, t= 2.65, *p*=.0081 and DLPFC, t= 2.401, *p*=.0016; PMA > SMA, t= 3.414, *p*=.0007 and DLPFC, t= 3.167, *p*=.0015). Consistent with trends for reach path magnitude, the effects of stimulation on reach path deviation were significantly greater in the younger relative to the older group (*p*<0.001 and *p*<0.001, respectively), but no significant region by group interaction was observed. As with reach path magnitude, there was also a significant effect of target at both time points (both *p*<0.001) and target by group interactions (*p*<0.001 and *p*<0.001, respectively). However, reverse trends were observed such that increased demands to counter the effects of gravity at successively higher target locations *increased* reach path deviation (UT>MT>LT) with the younger group exhibiting greater deviations, regardless of stimulation condition.

**Figure 5.**
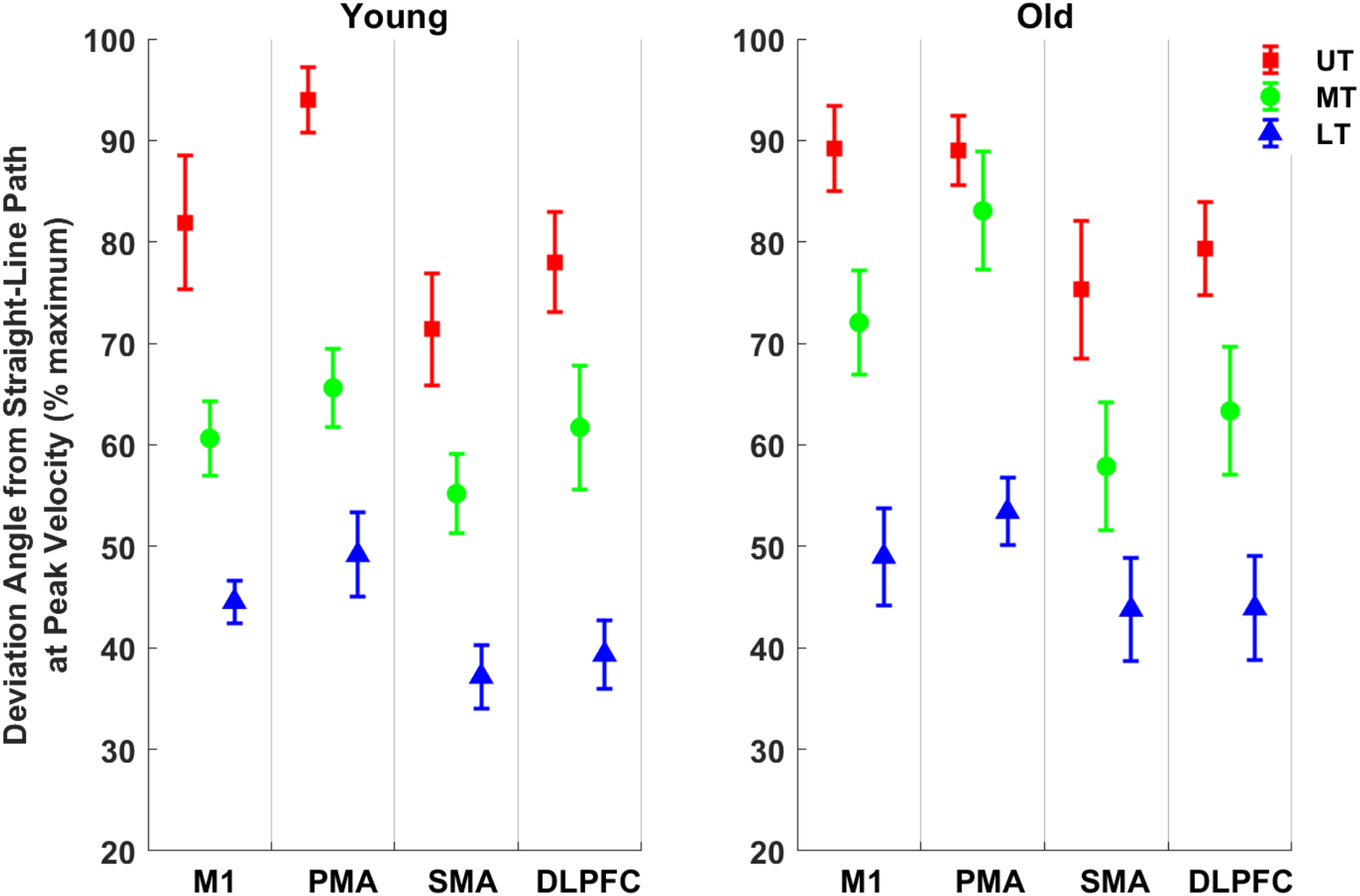
Trends in the regional effects of cortical stimulation on reach path deviation at the time of peak velocity in younger (left column) and older (right column) subjects at each target location under the 250-ms stimulation condition. Error bars represent standard error. Reach deviations across trials for all regions were normalized to the maximum deviation from a given subject.

**Figure 6.**
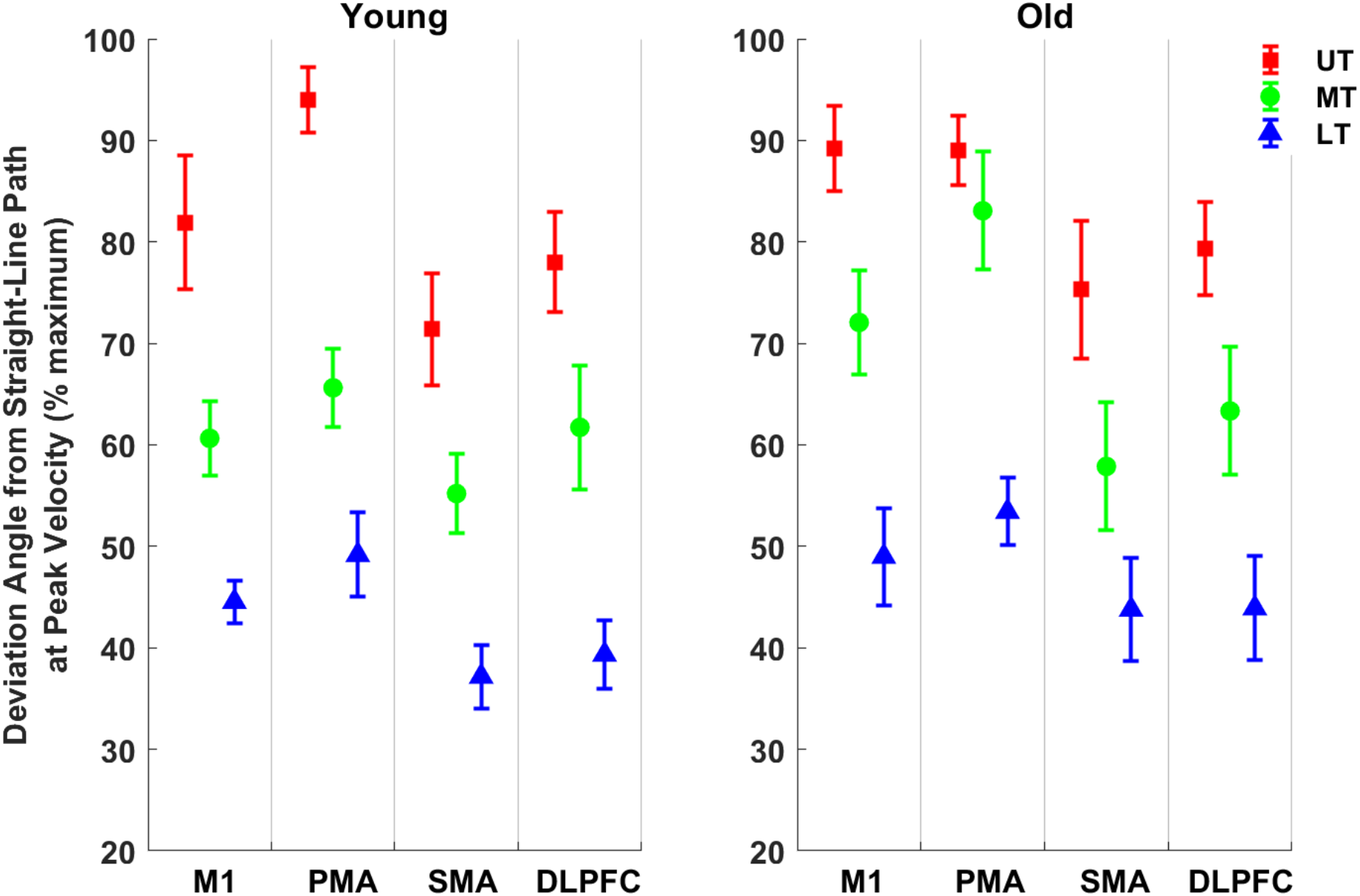
Trends in the regional effects of cortical stimulation on reach path deviation 50 ms after the time of peak velocity in younger (left column) and older (right column) subjects at each target location under the 250-ms stimulation condition. Error bars represent standard error. Reach deviations across trials for all regions were normalized to the maximum deviation from a given subject.

**Figure 7.**
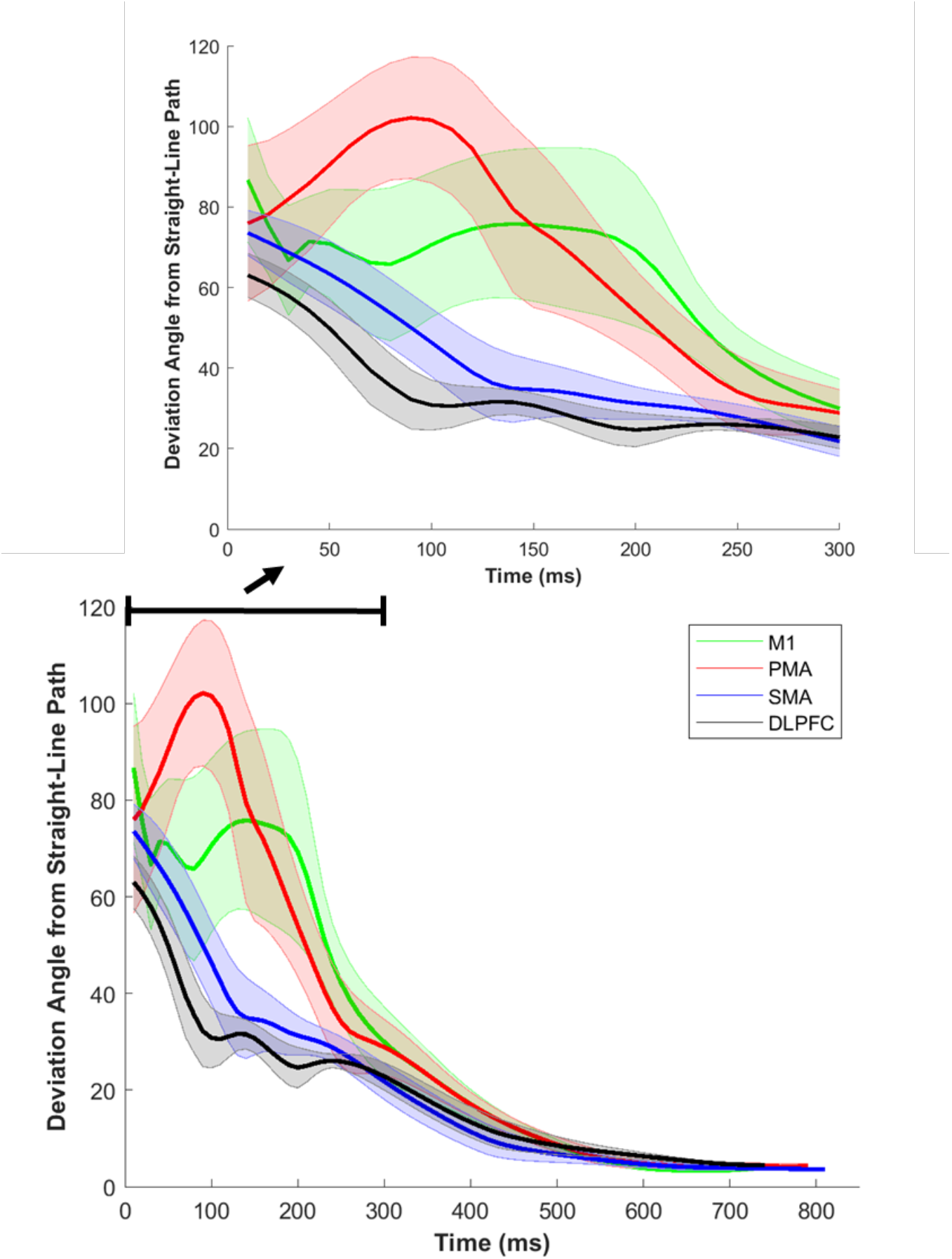
Exemplar reach path deviation from a single subject under the 250-ms stimulation condition. The bottom plot illustrates deviation of the reach vector from a straight-line path over the entire reach, whereas, the top plot depicts only the first 300 ms of the reach to more clearly show regional differences detected late in the movement trajectory at the time of peak velocity and 50 ms thereafter. Traces represent the mean of all trials when stimulation was delivered to a particular region, and shaded areas correspond to standard error.

Regional effects were also detected for overall reach path curvature (F_2,89_) =15.66, *p*<0.001, Fig 8). Exemplar reach path curvatures with stimulation to each cortical region are shown in Figure 9. Also similar to preliminary and final models on reach path deviation, M1 and PMA stimulation produced comparable increased curvature that was greater than SMA (M1, t= 2.988, *p*=.0028; PMA, t= 4.005, *p*<.001) and DLPFC (M1, t= 3.59, *p*=.0003; PMA, t= 4.613, *p*<.0001) stimulation. Reach path curvature was greater in the younger group (t= 3.68, *p*=.0002). Although a region by group interaction was not detected, there were region by target and group by target interactions (*p*=0.01 and *p*=0.008).

**Figure 8.**
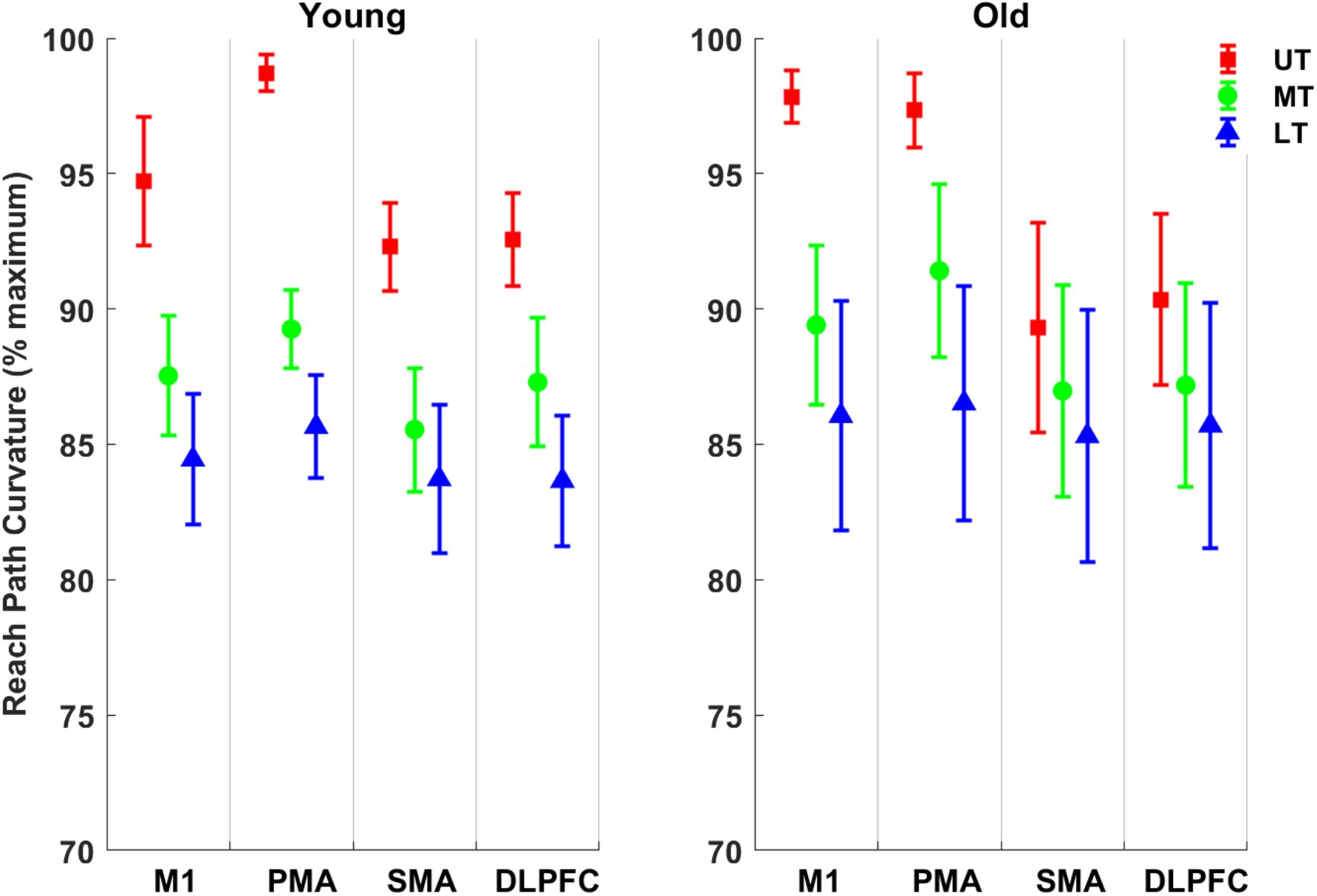
Trends in the regional effects of cortical stimulation on reach path curvature in younger (left column) and older (right column) subjects at each target location under the 250-ms stimulation condition. Error bars represent standard error. Reach curvature across trials for all regions were normalized to the maximum deviation from a given subject.

**Figure 9.**
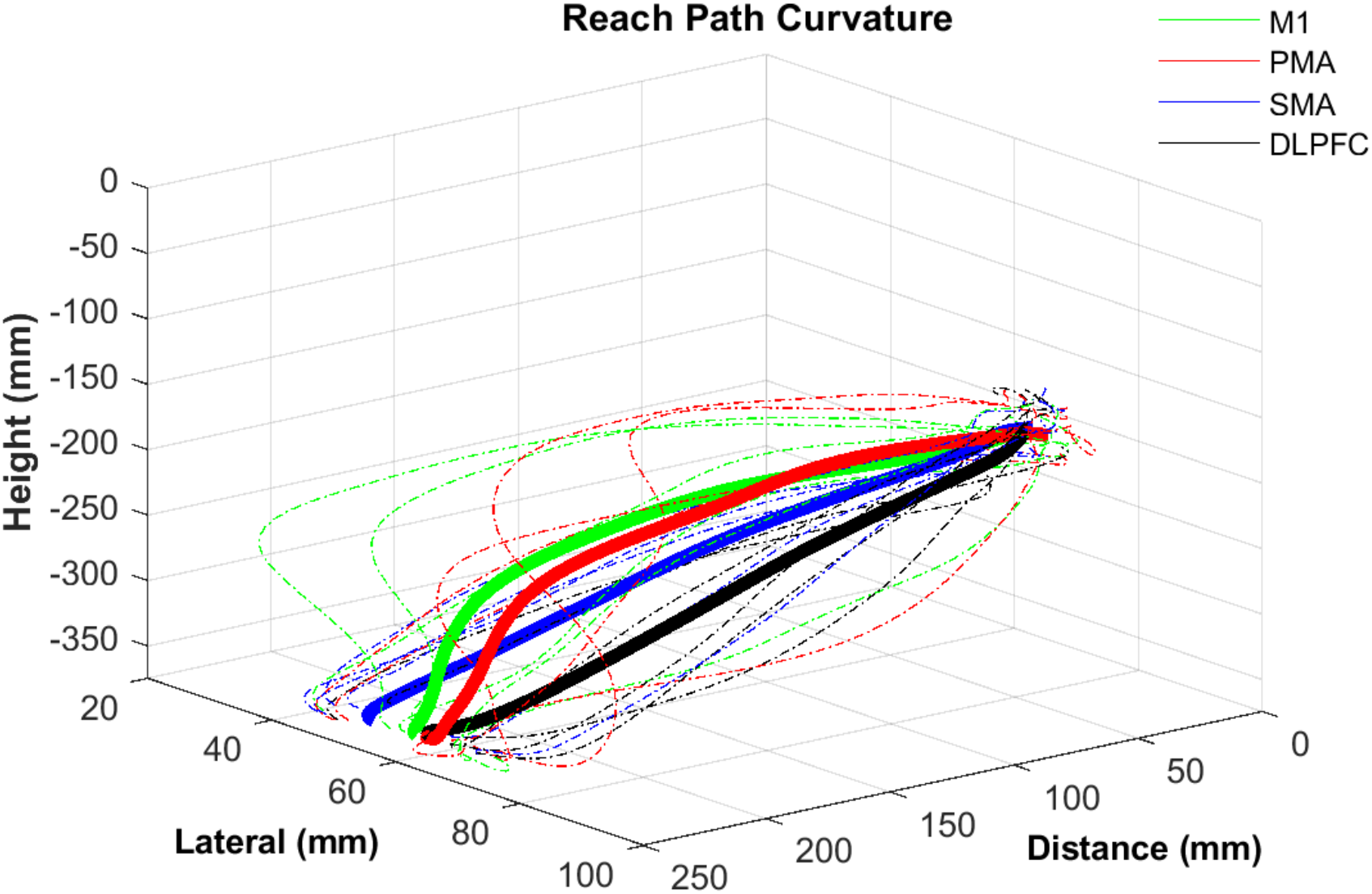
Exemplar reach path curvature from a single subject under the 250-ms stimulation condition. Bold traces represent the mean of all trials when stimulation was delivered to a particular region, and light traces correspond to standard error.

Separate models were used to examine the effects of stimulation on peak velocity of the reach and on the overall time-course of the movement trajectory. Mixed model analysis of peak velocity showed that stimulation increased peak velocity (F_2,89_) =59.28, *p*<0.001). There were also effects of group (*p*<0.001) and target (*p*<0.001), with older subjects exhibiting *higher* peak velocities relative to younger subjects and lower peak velocities at successively higher targets (UT<MT<LT). Mixed model analysis of the time-course of the movement revealed an effect of stimulation on the time that elapsed before (F_2,89_) =3.75, *p*<0.001) and after (F_2,89_) =1.95, *p*<0.001) peak velocity (Fig 10). Cortical stimulation reduced the rise time to peak velocity but increased the time that elapsed after peak velocity. Although stimulation produced stronger effects on the amplitude of reach kinematics in younger subjects, this group also exhibited less time before (*p*<0.001) and after peak velocity (*p*<0.001) despite reaching lower peak velocities.

**Figure 10.**
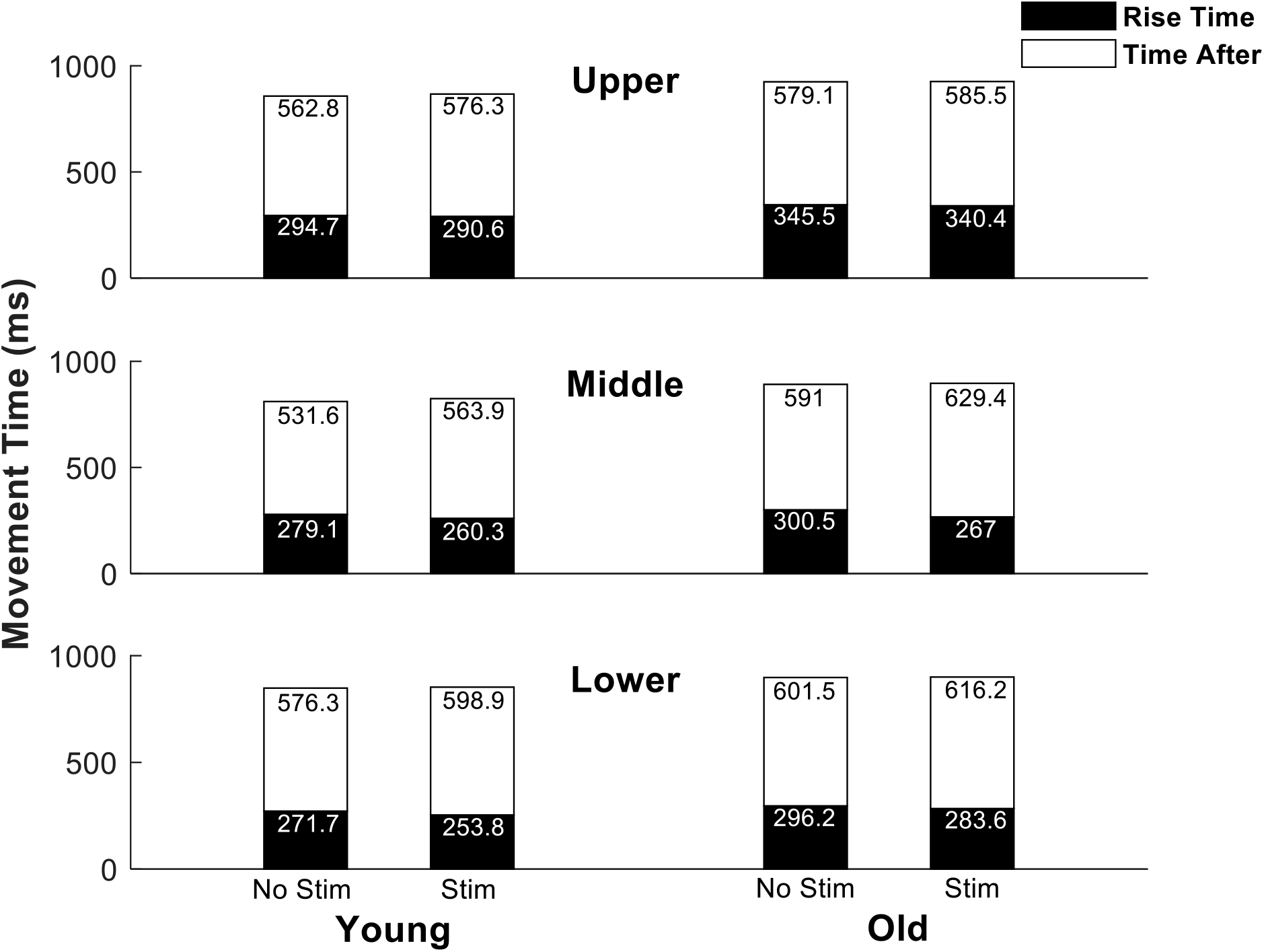
Figure 10. Stacked bar plot showing rise time and time after peak velocity under no-stimulation and 250-ms stimulation conditions. Note the decreased rise time and increased time after peak velocity with stimulation. Also note slower rise time and time after peak velocity in old vs. young groups.

Acknowledging that cortical stimulation was at or above rMT to three of the four brain regions, EMG recordings were inspected to determine the incidence of MEPs in the peristimulus time window (10-30 ms after stimulation onset). Overall, MEPs in shoulder and elbow muscles were evident in less than half of all cases for all muscles (Fig. 11). M1 stimulation produced a slightly higher overall incidence of MEPs (33.9%) relative to PMA stimulation (29.1%). Stimulation to both regions generated higher rates of MEPs relative to SMA (16.9%) and DLPFC (0%) stimulation. M1 stimulation seemed to recruit shoulder muscles to a greater extent than PMA stimulation (32.4% and 19.4%, respectively) but the inverse pattern was observed to a lesser extent for muscles crossing the elbow (35.3% and 39.2%, respectively. Distinct patterns in the incidence of deltoid *MEPs between M1 and PMA* remained fairly constant across target locations (UT: 30.3% vs 21.5%; MT: 36.6% vs 18.4%; LT: 30.3% vs. 18.4%). There was some degree of consistency in the pattern for triceps MEPs, but there was a distinct pattern in occurrence of biceps MEPs at successively higher target locations (LT: 29.4% vs. 23.5%, MT: <1% vs. 29.4% UT: <1% vs. 41.2%).

**Figure 11.**
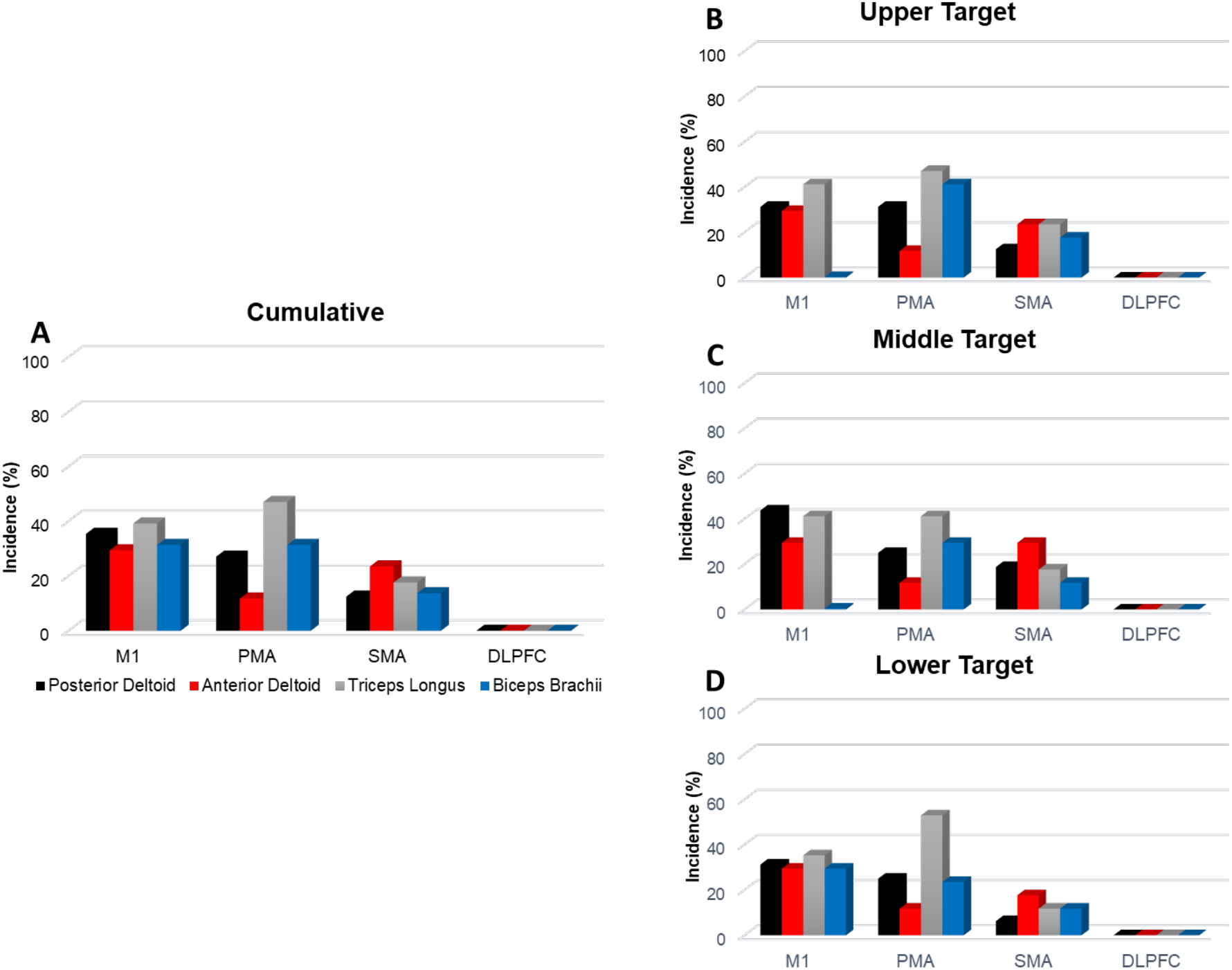
MEP incidence from stimulation to each brain region **A)** overall and for **B)** upper **C)** middle and **D)** lower target locations. Colored bars correspond to posterior deltoid (black), anterior deltoid (red), long head of the triceps (grey), and biceps brachii (blue).

## Discussion

### Summary

Consistent with prior work,^28,31^ differences in reach profiles were observed between age groups, with notably shorter reaction times evident in the younger group. Stimulation pulses to any brain region 50 ms before and 100 ms after the onset of the visual cue shorted reaction times, complicating the analysis of regional effects. While some of the early effects on the reach path could have been due to differences in reaction time, regional effects were detected, with PMA stimulation increasing early movement magnitude and later deviation from a straight-line path. PMA stimulation produced comparable effects to M1 stimulation, which served as a positive control because of its relative position downstream in the final common pathway for limb control.

The comparison of no stimulation vs. 250 ms post-go cue stimulation avoided the confound of differing reaction times. In this comparison, PMA stimulation continued to increase early reach path magnitude, with lesser effects of M1, SMA, and DLPFC stimulation. Early reach magnitude was also decreased by the height of the target, indicating an effect of antigravity movement requirements. Despite the lack of any effects of stimulation on reach path deviation early in the movement trajectory, there were effects later in the movement, after peak velocity and during target homing-in. PMA and M1 stimulation did not influence reach path magnitude around the time of peak velocity and 50 ms thereafter, but stimulation at both locations increased deviation from the straight-line path to an extent that was similar but greater than SMA and DLPFC stimulation. Opposite from trends in reach path magnitude, reach path deviation *increased* by target height. Overall path curvature was increased to a greater extent with PMA and M1 stimulation relative to SMA and DLPFC stimulation. Although stimulation, in any region, had milder effects on the reaches of the older subjects, this group exhibited higher peak velocity, yet increased time, before and after peak velocity relative to the younger group. We interpret these preliminary findings as support for a role of PMA in sensory-guided movement that is reduced by age.

### Reaction time effects

TMS interacts with human research participants through a few modalities, the pulsed magnetic field induced in the brain being just one of them. The solenoid-like vibration of the coil causes a clicking sound and tactile sensation. Induced electric currents in the scalp cause significant stimulation of sensory and motor nerves. These non-specific peripheral effects lead to brain input through cranial and cervical spinal nerves, affecting some central sensorimotor processing.^46^ It is therefore important to first understand reaction-time effects and also distinguish whether stimulation effects are regionally specific. Stimulation did have a non-regionally specific effect on reaction time, with stimulation just before and just after the go cue shortening the reaction time. Although participants were expecting a go cue, there were random delays built into its appearance, so the motor preparatory process should not have been progressing toward movement at any specific time, and there were catch trials in which no stimulation was given.

It has been demonstrated that auditory cues lead to faster reaction times than visual cues^47^ so it is unsurprising that an auditory cue occurring even after a visual cue could shorten reaction time. The sound produced by discharge of current into the coil was not intended as a cue. However, since reaction times were shortened by more than would be expected from the auditory-visual cue effect (about 30 ms), it is likely that there was some functional cueing effect of TMS. While the TMS pulses should not have elicited startle-like reactions, given that they occurred frequently, where relatively low loudness, and with explanation, coil discharge seems to have an indirect effect on reaction time.

### Effects on the early impulse control phase

The initial phase of a reaching movement can be characterized as a ballistic, approximately straight-line trajectory towards the target. Stimulation could interfere with planning and execution of movement in two main ways: a) increasing the level of activation in neurons that are already activated and b) inhibiting neurons over a longer period, reducing prolonged activity. These effects may be negligible because by the time upper motor neurons activate lower motor neurons, muscles contract and to generate movement.

The fact that PMA stimulation increased early reach path magnitude with no effect of stimulation timing suggests a role for PMA as a brake on movement during the time-range studied, a role consistent with known effects in action selection and important to investigate^48^. This is based on the argument that, without a strong time dependence of stimulation, local inhibitory effects would have dominated any lasting effects of stimulation. Increased curvature and deviation from straight-line path are consistent with the role of PMA in online control of movement.

PMA, M1, and SMA stimulation increased reach path magnitude within 100 ms of movement onset to a greater extent in the younger group. Reaching against gravity reduced movement amplitude particularly in old individuals, but there was no interaction to suggest an increased level of activity in antigravity motor representations that could be accessed by TMS. The effect of rTMS in this study was to increase forward movement regardless of stimulation target. It is possible that other regions, such as the reticular formation, play more of a role in the postural aspect of unsupported reaching^49^ and this general effect was more related to activation of the reticular formation.

### Effects on the later limb-target control phase

The later movement effects of stimulation over non-primary motor areas, measured at the time of peak velocity (i.e., after peak acceleration) reduced movement efficiency. There was no uniform pattern to the deviation, implying that inhibition of activity in these areas reduced the corrections to limb trajectory introduced in the initial, ballistic phase. Alternatively, the increased reach path observed early was compensated for later, but the compensation was insufficient to restore accuracy to the movement. In both interpretation, the role of PMA in homing-in to a target is suggested, and consistent with the two-component theory of goal-directed aiming^50^, and an age-related degradation in that behavior that is *not* related to PMA’s role^51,52^. PMA stimulation tended to have the strongest effect on deviation of the reach path in most subjects, despite a lack of statistical difference between PMA and M1 stimulation.

### Gravity effects and compensation

An initial hypothesis was that there would be specific regional control of the antigravity components of reaching movements. In particular, SMA seemed a likely node in the network for antigravity control, serving a role in production of antigravity postural adjustiments^53,54^. While target height affected kinematics, there was no specificity with regard to regional effects of stimulation. The lack of regional effects might suggests that the antigravity and vertical aspects of the movement shared a common cortical control mechanism. A special feature of reaching without antigravity support is the role of elbow flexors in every phase of the reach. Despite the fact that the elbow eventually extends, the elbow flexors lift the forearm off the support surface, and act eccentrically to control gravity-actuated forward extension and then support the forearm at the end-point of the reach. This is consistent with minimization in co-contraction and energy use in practiced movements.

More curved reach paths were exhibited at successively higher targets, which might reflect a compensatory strategy to offset the increased magnitude early and deviation late in the movement trajectory while also countering the effects of gravity. If so, such a compensation might require increased time after peak velocity to allow for online control processes to hone the reach vector into the target.

### Age effects

Despite faster reaction times, younger subjects also showed more sensitivity to stimulation effects both early in the impulse control phase (i.e., reach path magnitude), later in the limb-target control phase (i.e., reach path deviation), and throughout the entire movement (i.e., reach path curvature). This might be explained by one or more factors. The temporal pattern of reach-related activity could be more spread out in older individuals, reducing the effect of interference at any particular time. Cortical processing is reduced in older individual which may be due to an age-related decline in the integrity of cortical circuits. It is possible, therefore, that stimulation did not engage circuitry as robustly as in younger individuals. As part of preliminary analyses from these experiments, we found a higher amount of co-activation in the axial muscles of older subjects. The resulting joint stiffness may have reduced brief changes in neuronal activity elicited by stimulation.

Also despite more perturbation of initial reach movement trajectories, younger subjects were more readily able to coordinate the time-course of the movement trajectory and process sensory feedback more efficiently to enable online control of the reach.

### Limitations

One limitation of this study is sample size, both in number of participants and number of reaches. The latter was restricted due to concern about fatigue but could have been handled by multiple sessions or fewer trial types. With further refinement of the experimental paradigm used in this study, it is reasonable to test a larger number of participants over a broader age range, and in a number of neurological diagnoses, starting with stroke.

Another limitation relates to the use of a passive motion capture system. Line-of-sight was an issue, particularly as the participants were surrounded by equipment and an experimenter. This necessitated some gap filling when reflective markers were lost to view. This had little practical effect on the data. Another limitation of the study was the lack of neuronavigation-guided stimulation. The scalp locations and coil orientations for brain stimulation were selected by surface anatomy and relationships with optimal sites for two muscles. While we would have preferred to use stereotactic neuronavigation, this would have been in concert with such functional anatomy in order to confirm distance from the primary motor cortex. Future work should investigate use of MRI functional and anatomical markers for non-primary motor areas and control areas.

### Confounds

MEPs were elicited on some trials and broadly represent the interaction of stimulation with a cortical motor system that is not at rest. The incidence of MEP was dependent on target height, but different for M1 and PMA. A greater demand to overcome gravity at higher target locations could explain the increased incidence of MEPs for PMA stimulation, but not the reverse effect for M1.

Localization of TMS current is subject to the size and shape of the induced magnetic field. While several square centimeters of cortical surface receive significant currents, the effects are more restricted due to threshold effects^55^. So while TMS over M1, PMA, and SMA may affect neighboring areas – and in the case of SMA, the contralateral homolog – those effects are not likely to significantly affect neuronal activity. But SMA stimulation has other limitations, in terms of its distance from the scalp and variability in location^56^.

This was an *undesired* effect of the TMS paradigm used in this study. It is not possible to tell if the MEP was directly evoked from M1 regardless of coil position due to spread of induced current from non-M1 locations to M1 vs. through activation of the region to which the stimulation was directed. The mechanism of TMS pulse train interference depends on depolarizing current entering neurons; the effects include excitation and recurrent inhibition of various neuronal elements. Ideally, MEP occurrence would have been reduced further, but the procedure for doing so was impractical with the set-up used. As a first investigation into unconstrained reaching, we also wanted to ensure the stimulation would have measurable effects and, therefore, opted for stronger as opposed to weaker stimulation.

The idea that regional effects were driven, at least in part, by inhibition within these cortical areas was drawn from the literature using the same, or similar, technique. This interpretation is consistent with our observation that PMA stimulation increased movement amplitude in a manner that rivaled or exceeded M1 stimulation, the latter of which was intended to serve as a positive control given its predominant role in movement execution. And the relatively low time dependence of effects is more consistent with a longer-lasting inhibition than with selective increased activation in dynamically activated subsets of neurons.

### Future Directions

The primary objective of this study was to demonstrate the disruptive effects induced by brief trains of TMS in exploring the functional role of cortical regions in controlling unconstrained reaches against gravity. A goal for this study is to inform future work examining the role of these brain regions on motor control after stroke to explain and facilitate the recovery of arm function. Before addressing this goal directly, it will be important to refine the method further. For example, it has been shown that the timing of disruption can be achieved with single stimuli In addition, titrating stimulation intensities may be useful in distinguishing between excitatory and inhibitory effects.

## Conclusion

The study was an exploration of the effects of brief trains of TMS, used to interfere with regional cortical brain function, on unconstrained reaching with a vertical component. We found that TMS over M1 and (dorsal) PMA caused increased magnitude and deviation of the early phase of reaching that did not depend on the timing of the stimulation after the Go cue. This, and other results, provide an ongoing role of PMA in visually guided movement *after* movement initiation. For older subjects, TMS effects were weaker in the early reach but led to more disruption in later, suggesting an age-related reduction in sensorimotor processing flexibility for online control of unconstrained reaching.

## Author contributions statement

M.A.U. analyzed data and drafted the manuscript. J.T. analyzed data and wrote parts of the manuscript. G.P.M. did statistical analysis and writing. N.K. and L.W. had a primary role in data collection and protocol development. O.L., S.P.S., I.J, contributed to design and interpretation of the experiments. G.F.W. conceived of the project and was involved in every aspect of it. All authors approved the final version of the manuscript.

## Acknowledgments

Data collection was performed while GFW was a Senior Fellow at KU Leuven, on sabbatical from the University of Maryland, Baltimore and on Extended Educational Leave from the Dept. of Veteran Affairs. We would like to thank Rob Meugens for construction and initial programming of the reaching target system. Lianne Zevenbergen & Mariska Wesseling provided MATLAB scripts and advice for EMG and kinematic processing, respectively.

